# Crk mediates Csk-Hippo signaling independently of Yap tyrosine phosphorylation to induce cell extrusion

**DOI:** 10.1101/2024.06.27.601065

**Authors:** Abdul Hannan, Qian Wang, Yihua Wu, Neoklis Makrides, Xiuxia Qu, Junhao Mao, Jianwen Que, Wellington Cardoso, Xin Zhang

## Abstract

Src family kinases (SFKs), including Src, Fyn and Yes, play important roles in development and cancer. Despite being first discovered as the <u>Y</u>es-<u>a</u>ssociated <u>p</u>rotein, the regulation of Yap by SFKs remains poorly understood. Here, through single-cell analysis and genetic lineage tracing, we show that the pan-epithelial ablation of C-terminal Src kinase (Csk) in the lacrimal gland unleashes broad Src signaling but specifically causes extrusion and apoptosis of acinar progenitors at a time when they are shielded by myoepithelial cells from the basement membrane. *Csk* mutants can be phenocopied by constitutively active *Yap* and rescued by deleting *Yap* or *Taz*, indicating a significant functional overlap between Src and Yap signaling. Although Src-induced tyrosine phosphorylation has long been believed to regulate Yap activity, we find that mutating these tyrosine residues in both *Yap* and *Taz* fails to perturb mouse development or alleviate the *Csk* lacrimal gland phenotype. In contrast, Yap loses Hippo signaling-dependent serine phosphorylation and translocates into the nucleus in *Csk* mutants. Further chemical genetics studies demonstrate that acute inhibition of Csk enhances Crk/CrkL phosphorylation and Rac1 activity, whereas removing *Crk*/*CrkL* or *Rac1*/*Rap1* ameliorates the *Csk* mutant phenotype. These results show that Src controls Hippo-Yap signaling through the Crk/CrkL-Rac/Rap axis to promote cell extrusion.

## Introduction

SFKs are among the earliest identified and most studied oncoproteins, playing important roles in tumor progression and metastasis ^1^. Numerous Src substrates have been identified, including cell cycle regulators, adhesion molecules and transcription factors, contributing to cell proliferation, adhesion and motility. Src can be phosphorylated by the non-receptor tyrosine kinase Csk at the C-terminal tyrosine 527 residue, leading to conformational changes that suppress its kinase activity, whereas auto-phosphorylation by Src itself at the tyrosine 416 residue locks it into the active state ^2^. This negative regulation of Src by Csk is critical, as systemic knockout of *Csk* in mice causes neural tube defects and embryonic lethality as early as E9.5 ^3,4^. Interestingly, ubiquitous loss of *Csk* in *Drosophila* led to overproliferation and decreased cell death, whereas mosaic *Csk* inactivation resulted in epithelial extrusion and apoptosis ^5^. The same behavior in cell extrusion has been observed in isolated Src-transformed mammalian cells within normal epithelial sheets ^6^, yet the underlying mechanism is poorly understood.

Lacrimal gland development begins as a thickening of the epithelium at the superior conjunctival fornix, which subsequently invades the underlying mesenchyme to form a highly branched structure during embryogenesis ^7^. However, the cellular identity of the lacrimal gland is not fully established until after birth, when acinar cells coalesce into closely packed clusters infiltrated by myoepithelial cells and are connected by duct cells that ultimately reach the ocular surface ^8,9^. Many studies have focused on the embryonic development of the lacrimal gland ^7^. In particular, we and others have shown that FGF signaling interacts with multiple other pathways, including BMP, EGF, Notch and Wnt, to regulate lacrimal gland budding, elongation and branching morphogenesis ^10–20^. In contrast, relatively little is known about the maturation and homeostasis of the lacrimal gland during the postnatal stage.

In this study, we investigated the role of Csk in lacrimal gland development. We showed that an epithelial-specific knockout of *Csk* did not affect branching morphogenesis during embryogenesis, but the lacrimal gland degenerated postnatally with dilated lumens lacking acinar cells. This phenotype was caused by the apical extrusion of lacrimal gland progenitors, which could be reproduced by constitutive activation of *Yap* at a critical time window before acinar cells are exposed to the basement membrane and rescued by the deletion of either *Yap* or *Taz*. By mutating tyrosine phosphorylation sites in Yap and Taz, we demonstrated that Src does not directly regulate Yap/Taz through tyrosine phosphorylation as previously reported ^21–24^. Instead, *Csk* deficiency led to Crk/CrkL-mediated stimulation of Rac1 and Rap1 that promoted the nuclear localization of Yap, and the *Csk* mutant phenotype can be rescued by the deletion of *Crk/CrkL*, *Rac1* or *Rap1*. These results demonstrate that the Crk/CrkL-Rac1/Rap1-Yap/Taz axis is responsible for Src-induced cell extrusion.

## Results

### Genetic deletion of *Csk* during lacrimal gland development resulted in luminal expansion and acini loss

To explore the role of *Csk* in the lacrimal gland, we generated a conditional knockout of *Csk* using the *Le-Cre* driver, which is active in ocular surface progenitor cells during embryonic development ^13^. This led to the induction of the *R26R* Cre reporter in the entire lacrimal gland epithelium, but not in the mesenchyme (Fig. 1A). Interestingly, although *Le-Cre;Csk^f/f^* (*Csk^CKO^)* lacrimal glands did not display overt branching defects during embryonic development, their end buds appeared to be grossly dilated by P2 as shown by whole-mount Xgal or SMA/Krt14 staining (Fig. 1A, arrowheads). In sections, instead of forming the closely packed clusters as in the controls, the Xgal+ mutant epithelial cells encircled dilated lumens (supplementary Fig. 1A-B, arrows and arrowheads). By P21, the *Csk^CKO^*mutant lacrimal glands were clearly atrophic, interspersed with numerous melanocytes (Fig. 1A, dotted line and arrowheads).

**Figure 1.**
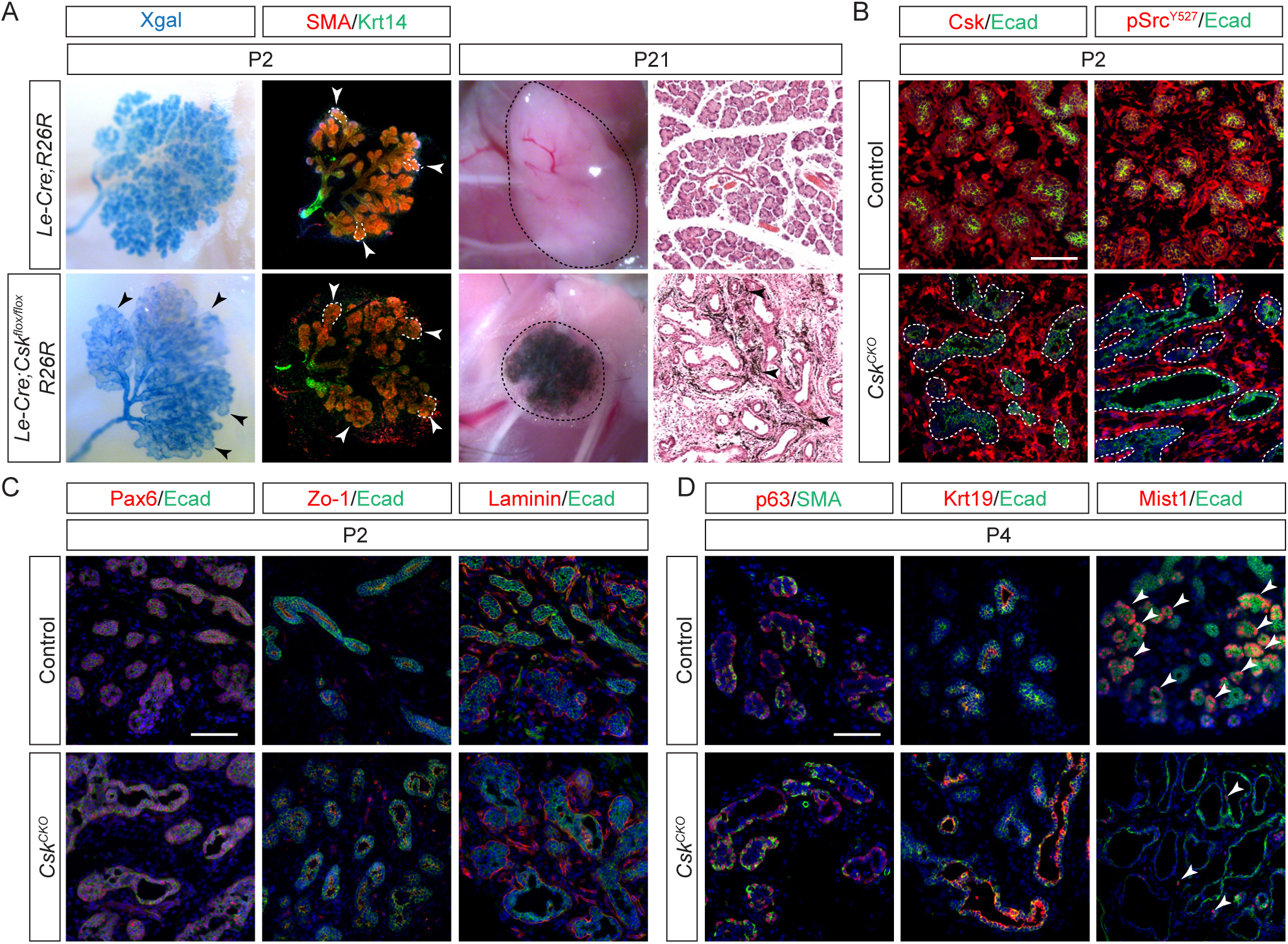
The epithelial-specific deletion of *Csk* disrupted postnatal lacrimal gland development. **(A)** Driven by the epithelium-specific *Le-Cre* as indicated by *R26R* reporter activity (Xgal staining), the conditional knockout of *Csk* resulted in the dilation of lacrimal gland end buds (black arrowheads) at P2, evident in whole-mount staining of SMA and Krt14 (white arrowheads and dotted outlines). At P21, the atrophic lacrimal gland (dotted lines) in *Le-Cre;Csk^flox/flox^* (*Csk^CKO^*) mutants contained numerous melanocytes (arrowheads). **(B)** Consistent with the role of Csk as the C-terminal kinase of Src, the loss of Csk expression in the expanded epithelial lumens was accompanied by the absence of Src-Y527 phosphorylation (pSrc^Y527^) (dotted lines). **(C)** *Csk^CKO^*mutants retained the lacrimal gland progenitor marker Pax6, E-caderin+ adherens junctions, ZO-1 tight junctions and Laminin+ basement membrane at P2. **(D)** In contrast to the persistent expression of myoepithelial markers p63 and SMA and ductal marker Krt19, there was a drastic reduction in the acinar marker Mist1 (arrowheads). Scale bars: 100 µm.

We next examined the molecular changes in *Csk* mutant lacrimal glands. In line with the specificity of the *Le-Cre* driver, Csk was selectively depleted in the E-cadherin-expressing epithelial cells, which also lost pSrc^Y527^ phosphorylation (Fig. 1B, dotted lines). This confirms that Csk is indeed the obligatory kinase for the Src Y527 residue. Pax6 is a progenitor marker for the lacrimal gland epithelium, flanked by the apical Zo-1 and the basal Laminin staining, whereas Col2a1 is restricted to the mesenchyme. In P2 *Csk^CKO^*mutants, these markers were found in their proper domains and the cell proliferation rate was also unchanged (Fig. 1C and supplementary Fig. 1C-G). By P4, however, although SMA/p63+ myoepithelial and Krt19+ ductal cells were observed around the dilated lumens, the acinar marker Mist1 was mostly absent in *Csk^CKO^* mutants (Fig. 1D, arrowheads). Therefore, while *Csk* is dispensable for the branching morphogenesis of the lacrimal gland during embryonic development, it is necessary for the postnatal maturation of the acinar ductal network.

### Single-cell RNAseq revealed epithelial cell extrusion and apoptosis in *Csk* mutant lacrimal glands

To understand the molecular mechanisms underlying the *Csk* mutant phenotype, we performed single-cell RNA sequencing (scRNAseq) of P1 lacrimal glands. To avoid batch effects, control and mutant lacrimal gland cells were pooled together in library preparation after labelling with Cell Multiplexing Oligos (CMOs), which allowed later bioinformatic deconvolution of the sequencing reads. We sequenced 4,178 control and 5,853 *Csk^CKO^* mutant cells at a depth of about 1,800 genes per cell (Fig. 2A). Following the Seurat workflow to reintegrate the single-cell profiles for unsupervised clustering, the major cell types were annotated in the Uniform Manifold Approximation and Projection (UMAP) plots using known lacrimal gland markers (supplementary Figs. 2 and 3) ^25,26^. Like a previous scRNAseq study, we observed that most neonatal lacrimal gland cells belonged to either the epithelium or mesenchyme, but the higher numbers of cells sequenced in our experiment permitted us to discern much greater heterogeneities in each population (Fig. 2B) ^27^. Surprisingly, the *Csk^CKO^* mutant was enriched in one of the mesenchymal clusters (Mes6) compared to the control (Fig. 2C), despite our conditional knockout of *Csk* being targeted to the epithelium. We thus performed differential expression gene (DEG) analysis of this population, which identified a top candidate *Cdkn2b* highly specific to this cluster (Fig. 2D). Next, we labeled the epithelial cell lineage in the *Csk^CKO^* mutant using the tdTomato-expressing *Ai9* Cre reporter and showed by RNAscope in situ hybridization that *Cdkn2b* expression was restricted to the tdTomato+ cells.

**Figure 2.**
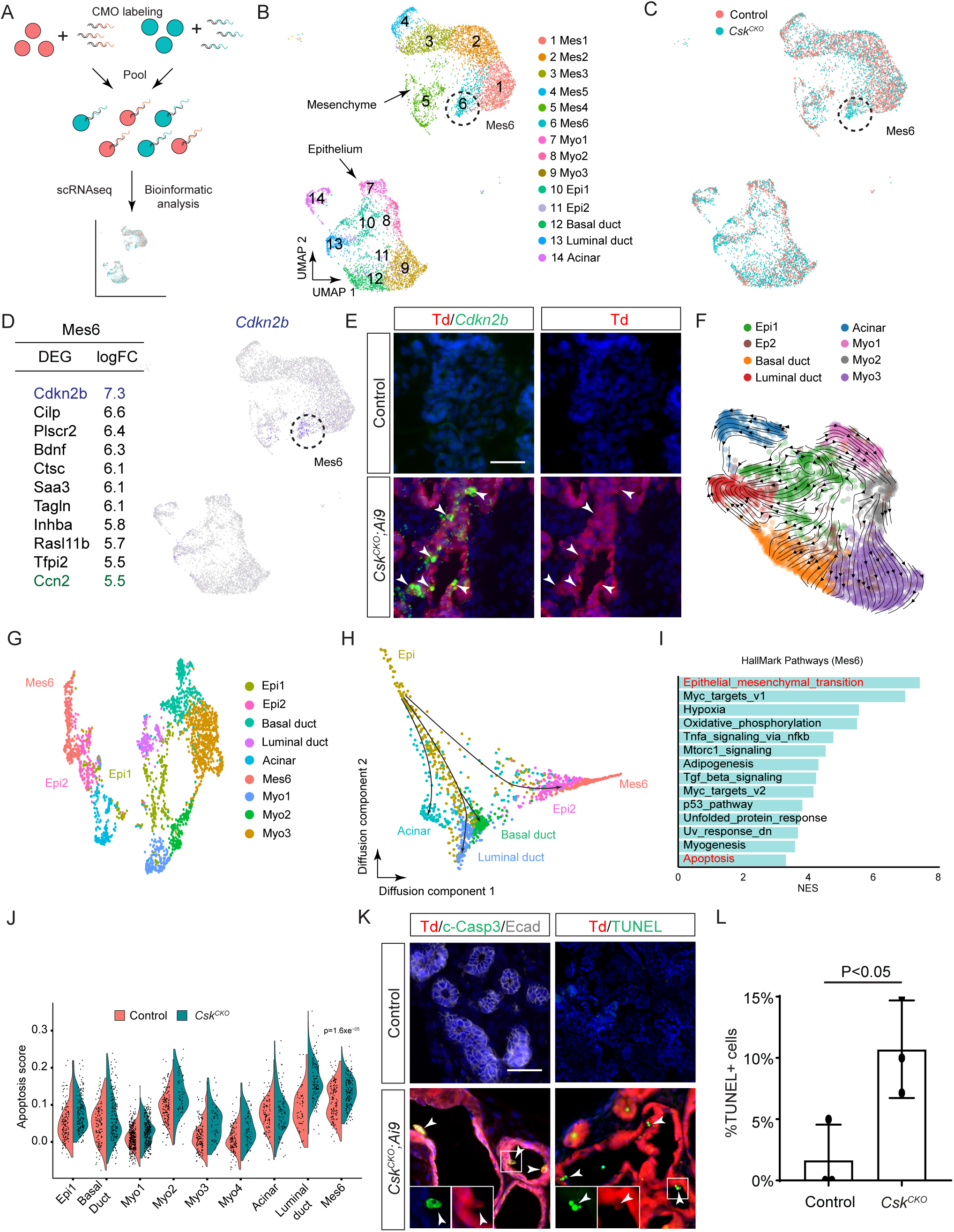
*Csk* mutant lacrimal glands exhibited epithelial cell extrusion and apoptosis. **(A)** Schematic diagram of the multiplex scRNAseq workflow to minimize batch effect. Cell suspensions from P1 control and *Csk^CKO^* mutant lacrimal gland cells were labeled with distinct CMO oligos, enabling demultiplexing by bioinformatic analysis after pooled scRNAseq. **(B)** The UMAP depiction of scRNAseq data. For clarity, only the epithelial and mesenchymal cell clusters were shown and the full UMAP clusters are presented in supplementary Figure 2B. **(C)** Although *Csk* was ablated only in the lacrimal gland epithelium, *Csk^CKO^*mutant cells were enriched in Mesenchymal cluster 6 (Mes6, circled). **(D)** Differential expression gene (DEG) analysis and feature plot showed that *Cdkn2b* expression was highly specific to the Mes6 cluster. **(E)** Using the *Ai9* reporter to follow *Le-Cre* expressing epithelial cells, RNAscope analysis indicated that *Cdkn2b* expression was restricted to the tdTomato-positive lacrimal gland epithelium (arrowheads), suggesting that Mes6 cluster cells were derived from the epithelial lineage. **(F)** The RNA velocity field revealed the developmental trajectories radiating from the Epi1 cluster toward other lacrimal gland epithelial cells, identifying the Epi1 cells as the progenitor population. **(G)** Re-clustering Mes6 cells with other epithelial cells showed that Mes6 cells are adjacent to a mixture of the Epi1 progenitors and Epi2 transitional cells. **(H)** Diffusion map showed the differentiation trajectories from the Epi1 progenitor cells to the acinar, myoepithelial, basal and luminal duct and Mes6 cells. **(I)** GSEA analysis of Mes6 cells highlighted the epithelial-mesenchymal transition and apoptosis pathways. **(J)** *Csk^CKO^*mutant Mes6 clusters exhibited elevated apoptotic signatures. **(K)** Cleaved caspase 3 (c-Casp3) and TUNEL staining confirmed increased apoptosis in *Csk^CKO^*mutants. Importantly, although these apoptotic cells retained the epithelial lineage marker Td, they have delaminated from the epithelium and extruded into the lumen (arrowheads). **(L)** Quantification of TUNEL+ cells in the lacrimal gland epithelium. Student’s t test, n=3, P<0.05. Scale bars: 200 µm.

**Figure 3.**
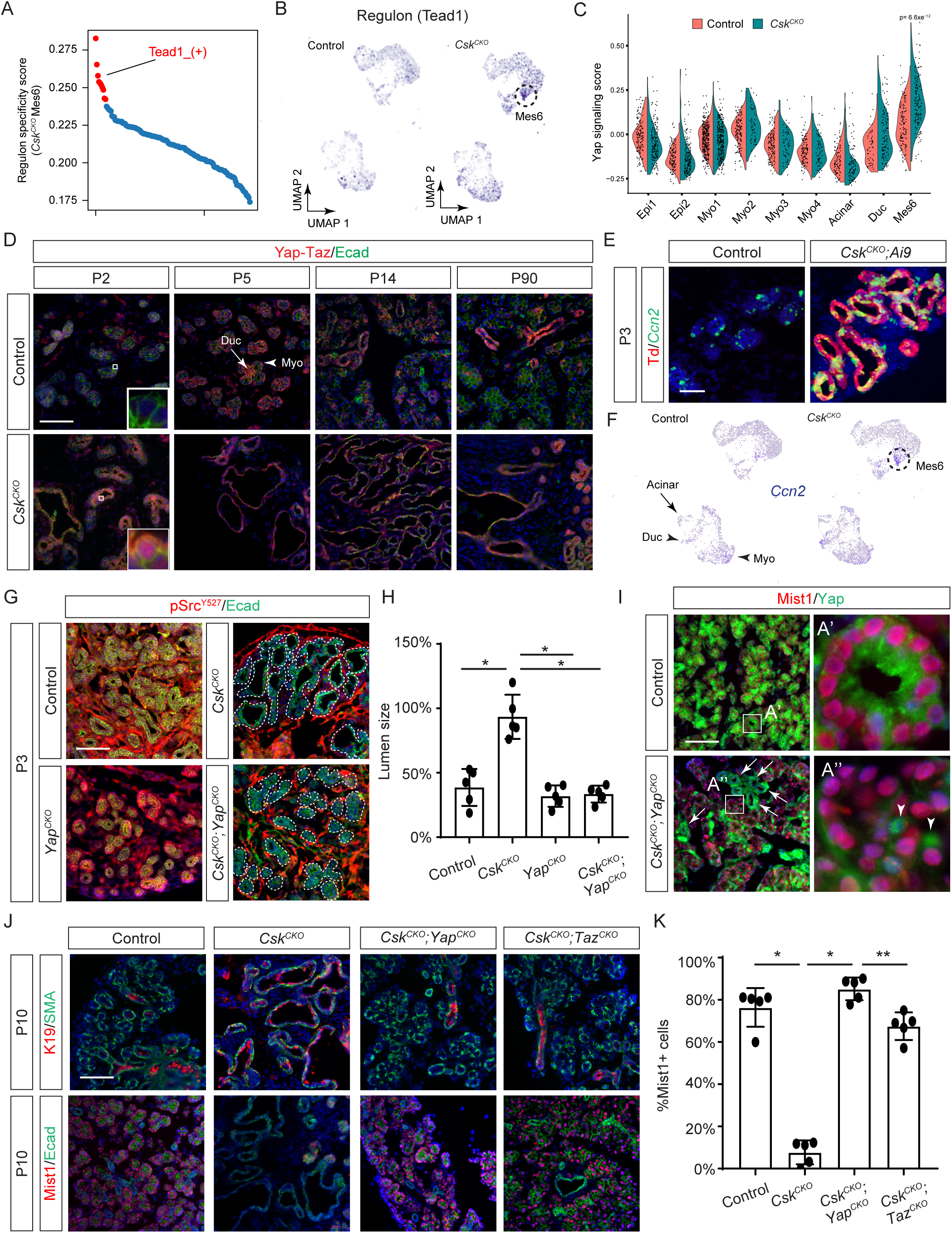
Yap and Taz deletion rescued *Csk* mutant phenotype. **(A)** SCENIC analysis identified that one of the top Mes6-specific regulons in *Csk^CKO^* mutant is activated by Tead1. **(B)** The feature plot shows the activation of the Tead1 regulon in *Csk^CKO^* mutant Mes6 cluster. **(C)** The Yap signaling signature in the Mes6 cluster was significantly upregulated in *Csk^CKO^* mutants. **(D)** In control lacrimal glands from P2 to P90, Yap/Taz staining was nuclear only in ductal (arrow) and myoepithelial (arrowhead) cells. In *Csk^CKO^* mutants, however, nuclear Yap/Taz was detected throughout the lacrimal gland epithelia. **(E)** *Ccn2* expression was restricted to Td-positive epithelial cells in *Csk^CKO^* mutants. **(F)** The Yap response gene *Ccn2* was highly expressed in the Mes6 cluster in *Csk^CKO^*mutants. **(G)** Although deletion of *Yap* didn’t reverse Src-Y527 phosphorylation, it rescued the lacrimal gland dilation phenotype in *Csk^CKO^*;*Yap^CKO^*mutants (dotted lines). **(H)** The lumen size was quantified as the ratio of the lumen area versus the overall epithelial area. One-way ANOVA, n=5, *P<0.0001. **(I)** The acinar marker Mist1 was recovered in *Csk^CKO^*;*Yap^CKO^*mutants, except in a few mosaic epithelial cells that retained Yap nuclear expression (arrows and arrowheads in A’ and A’’), indicating the cell autonomous function of Yap responsible for the *Csk* mutant phenotype. **(J)** Deletion of either *Yap* or *Taz* in *Csk^CKO^* mutants deflated Krt19 positive lumens and restored Mist1-expressing acini. **(K)** Quantification of Mist1+ cells. One-way ANOVA, n=5, *P<0.0001, **P<0.01. Scale bars: 100 µm.

This result suggested that, despite having a transcriptional profile characteristic of mesenchymal cells, the Mes6 cluster in the *Csk^CKO^* mutant was, in fact, derived from the lacrimal gland epithelium.

We conducted RNA velocity analysis to map the developmental lineages and cellular dynamics of the lacrimal gland. It identified the Epi1 cluster as the initial state of the epithelial lineage (supplementary Fig. 3B), which was evident in the velocity field where the central Epi1 vectors swirled in a pattern of self-renewal, while the peripheral ones radiated out toward myoepithelial, basal duct, luminal duct and acinar cells (Fig. 2F). Since our lineage tracing experiment demonstrated that the Mes6 cells were derived from the lacrimal gland epithelium, we next reclustered the Mes6 cells with other epithelial populations, which placed the Mes6 population adjacent to the Epi2 cluster mixed with some Epi1 cells (Fig. 2G). Accordingly, the diffusion map revealed a developmental trajectory from Epi1 traversing through the Epi2 state before reaching Mes6 (Fig. 2H), suggesting that the Mes6 cells were also derived from Epi1 progenitors as other lacrimal gland epithelial cell types ^28^. On the other hand, gene set enrichment analysis (GSEA) identified epithelial-mesenchymal-transition (EMT) as the most prominent pathway associated with the Mes6 cluster, explaining why Mes6 cells have acquired mesenchymal characteristics. Apoptosis was another pathway significantly enriched in the Mes6 population, with *Csk^CKO^* mutants displaying higher apoptosis scores than controls (Fig. 2I). This increase was confirmed by cleaved caspase-3 and TUNEL staining, which identified apoptotic cells adhering to the epithelial layers in *Csk^CKO^* mutants(Fig. 2K-L). Notably, these apoptotic cells were derived from the epithelial lineages, as evidenced by their continued expression of tdTomato (Fig. 2K, arrowheads), even though there was no apoptosis within the epithelial layer itself. These observations demonstrated that these cells were extruded apically from the epithelial layer into the lumen, where they subsequently underwent apoptosis.

### *Csk* mutant phenotype can be rescued by inactivation of Yap/Taz signaling

After uncovering a subset of lacrimal gland epithelial cells that underwent cell extrusion, we performed SCIENIC analysis to identify the transcriptional networks aberrantly activated by *Csk* ablation ^29^. One of the top Mes6-specific candidates was the Tead1 positive regulon, which was highly induced in Mes6 cells as shown by feature plots (Fig. 3A and B). The Tead family transcription factors are activated by the cofactors Yap and Taz, which undergo cytoplasmic to nuclear translocation in response to a variety of extracellular stimuli ^30–32^. Indeed, Yap signaling scores were most significantly increased in the Mes6 cells, but not in any other mesenchymal populations (Fig. 3C) ^33^. To confirm this finding, we examined the expression pattern of Yap/Taz proteins. In the control lacrimal gland, nuclear Yap-Taz staining was observed either in myoepithelial cells at the periphery of the epithelial cluster or in the duct cells at the center (Fig. 3D, arrow and arrowhead), but never in the acinar cells that made up the bulk of the E-cadherin+ epithelium (Fig. 3D, insert). In contrast, the epithelial lining of dilated lumens in *Csk^CKO^* mutants consistently displayed nuclear Yap-Taz staining from P2 to the adult stage. In line with this, the Yap response gene *Ccn2* (*Ctgf*) was sporadically expressed in the control lacrimal gland, but it was sharply upregulated in the tdTomato-labeled mutant epithelium (Fig. 3E). The ectopic *Ccn2* expression in *Csk^CKO^* mutants likely came from Mes6 cells since scRNAseq data showed that *Ccn2* was one of the top DEGs in this population (Fig. 2D) and specifically induced in the mutant gland (Fig. 3F). These results demonstrated that the loss of *Csk* led to the aberrant upregulation of Yap signaling in a subset of lacrimal gland epithelial cells.

The striking activation of Yap signaling in the lacrimal gland epithelium motivated us to ask whether it could account for the *Csk* mutant phenotype. To this end, we genetically ablated *Yap* in *Csk^CKO^* mutants, which did not affect the loss of pSrc^Y527^ phosphorylation, but suppressed the dilation of the epithelial lumen (Fig. 3G and H). This was paralleled by the recovery of Mist1 expression, indicating the rescue of acinar differentiation in *Csk^CKO^*;*Yap^CKO^* mutants (Fig. 3I).

Considering that Cre-mediated recombination is sometimes not fully penetrant, we searched for mutant cells that may have lost *Csk* but not *Yap*. Indeed, there were occasionally enlarged lumens in *Csk^CKO^*;*Yap^CKO^* mutants and even isolated Mist1+ cells within acinar clusters, but they inevitably retained nuclear Yap staining (Fig. 3I, arrows and arrowheads). These genetic mosaics demonstrated that nuclear Yap acts cell autonomously to disrupt acinar development in *Csk^CKO^* mutants.

Lastly, we investigated whether the genetic deletion of the Yap homolog Taz could also rescue the loss of Csk in lacrimal development. In striking contrast to dilated lumens in *Csk^CKO^* mutants, *Csk^CKO^*;*Taz^CKO^* mutants largely maintained a normal glandular architecture consisting of compact acinar clusters surrounded by SMA+ myoepithelial cells and connected by Krt19+ ductal networks (Fig. 3J). Similar to control and *Csk^CKO^*;*Yap^CKO^* animals, *Csk^CKO^*;*Taz^CKO^* mutants also displayed abundant expression of Mist1, although the percentage of Mist1+ cells was slightly reduced (Fig. 3J and K). Given the functional redundancy between Yap and Taz, these findings suggest that the overall increase in the Yap/Taz signaling dosage is responsible for the *Csk* mutant lacrimal gland phenotype.

### Yap activation targets the acinar progenitors unmoored from the basement membrane

How did Csk deletion cause such a profound lacrimal gland phenotype? We first considered whether this is due to defective cell differentiation by employing a series of tissue-specific and tamoxifen-inducible Cre drivers targeting specific cell lineages: *Mist1^Cre-ERT2^*for acinar cells, *SMA-Cre^ERT2^* for myoepithelial cells and *Krt7^Cre-ERT2^* for ductal cells (Fig. 4A) ^34–36^. These mutants were injected with Tamoxifen for three consecutive days from P1 to P3 and harvested at P6. The induction and specificity of each Cre driver were confirmed by either Ai9 or *R26^mTmG^* reporters, but none of these mutants exhibited any gross morphological defects (Fig. 4B).

**Figure 4.**
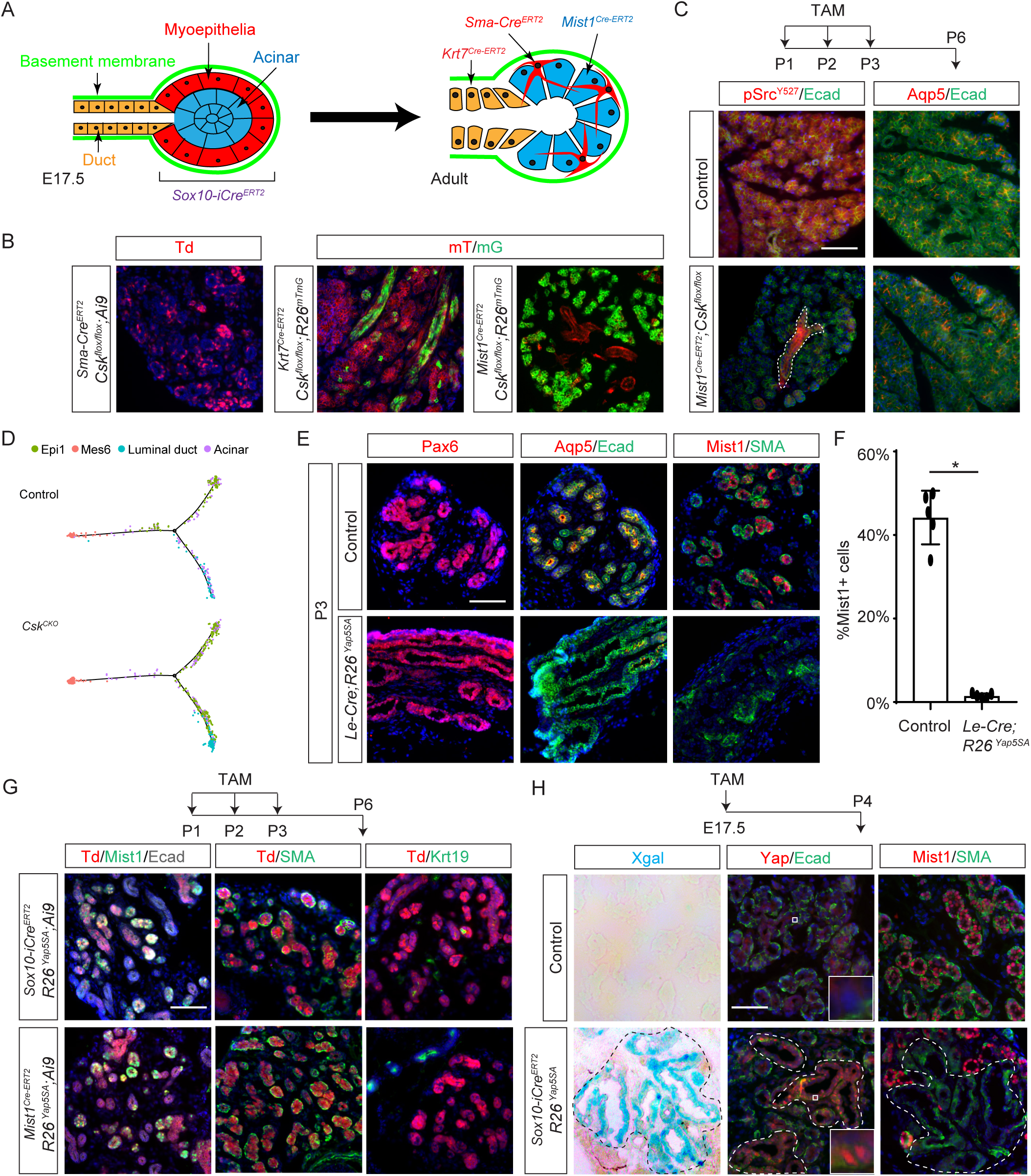
Yap activation specifically affects the acinar progenitors when they are separated from the basement membrane. **(A)** In the E17.5 lacrimal gland end bud targeted by *Sox10-iCre^ERT2^*, an inner mass of acinar progenitors is insulated by an outer layer of myoepithelial progenitor cells from exposure to the basement membrane. This seal is broken in the postnatal gland where the scattered myoepithelial cells can be targeted by *SMA-Cre^ERT2^*, acinar cells by *Mist1^Cre-ERT2^* and ductal cells by *Krt7^Cre-ERT2^* . **(B)** The cell-type-specific induction of *Mist1^Cre-ERT2^*, *Krt7^Cre-ERT2^* and *SMA-Cre^ERT2^*was confirmed by *Ai9* and *R26^mTmG^* fluorescent Cre reporters, but they did not cause a luminal dilation phenotype when crossed with the *Csk^flox/flox^* allele. **(C)** *Mist1^Cre-ERT2^*-mediated deletion of *Csk* led to the widespread loss of pSrc^Y527^ in acinar but not ductal cells (dotted line). Nevertheless, Aqp5 expression remained unchanged. **(D)** Pseudotime analysis of scRNAseq data suggested that *Csk* deletion disrupted the differentiation of the lacrimal gland progenitor cells (Epi1) but not the end stages of Mes6, acinar and ductal cells. **(E)** Induction of constitutively active Yap^5SA^ in lacrimal gland primordia by *Le-Cre* led to the dilation of epithelial lumens and loss of Aqp5+/Mist1+ acinar cells without affecting Pax6 and SMA expression. **(F)** Quantification of Mist1+ cells. Student’s t test, n=5, *P<0.0001. **(G)** Postnatal induction of the *Yap^5SA^* transgene by *Sox10-iCre^ERT2^* in lacrimal gland epithelial cells or by *Mist1^Cre-ERT2^*in acinar cells as shown by the Td reporter failed to perturb lacrimal gland morphology and the expression of the acinar marker Mist1, myoepithelial marker SMA and ductal marker Krt19. **(H)** Tamoxifen induction of *Sox10-iCre^ERT2^*;*Yap^5SA^*animals at E17.5 led to the ectopic activation of the *Yap^5SA^* transgene as indicated by the expression of nuclear localized Yap protein and LacZ reporter activity revealed by Xgal staining, resulting in luminal dilation and loss of Mist1 expression. Scale bars: 100 µm.

Notably, *Mist1^Cre-ERT2^*;*Csk^flox/flox^* mutants, while exhibiting an extensive loss of pSrc^Y527^ staining except in ductal cells (Fig. 4C, dotted line), still retained Aqp5-expressing acinar cells. These findings suggest that Csk function may not be essential for terminal differentiation within specific lacrimal gland cell lineages.

To understand why postnatal deletion of *Csk* failed to disrupt lacrimal gland development, we revisited the single-cell RNAseq data. Using Monocle to order single cells along pseudotime trajectories, we noticed that the Epi1 cluster displayed the most prominent displacement in *Csk* mutants, suggesting that the primary defect may lie within the epithelial progenitors (Fig. 4D) ^37^. To test this model, we took advantage of the *R26^Yap5SA^* allele, which can be induced by Cre to express constitutively active Yap^5SA^ protein resistant to suppression by Hippo signaling ^38^. In *Le-Cre*;*R26^Yap5SA^* mutant lacrimal gland at P3, Pax6 and E-cadherin staining revealed dilated lumens (Fig. 4E). Although SMA+ myoepithelial cells were present, the acinar markers Aqp5 and Mist1 were lost, demonstrating that activated Yap can recapitulate the *Csk^CKO^* mutant phenotype (Fig. 4E and F).

The lacrimal gland primordia consists of a Sox10^-^ stalk that gives rise to the duct and a Sox10^+^ bud that differentiates into myoepithelial and acinar cells (Fig. 4A). Genetic lineage tracing has revealed that myoepithelial cells are specified by E17.5, creating a protective cap over the acinar progenitors within the bud ^27,39^. This effectively insulates the acinar progenitors from the surrounding basement membrane until the myoepithelial cells begin to disperse after birth. We hypothesize that this transient separation from the basement membrane during the prenatal stage distinguishes acinar progenitors from the duct and myoepithelial cells, making them particularly prone to cell extrusion due to their lack of anchorage to the basement membrane. To test this hypothesis, we first administered Tamoxifen to *Sox10-iCre^ERT2^*;*R26^Yap5SA^*;*Ai9* and *Mist1^Cre-ERT2^*;*R26^Yap5SA^*;*Ai9* animals from P1 to P3 and analyzed them at P6 ^35,40^. While tdTomato expression confirmed efficient induction of *Sox10-iCre^ERT2^*and *Mist1^Cre-ERT2^* in the lacrimal gland epithelium and acinar cells, respectively, the expression of differentiation markers such as Mist1, SMA and Krt19 remained unchanged, indicating that Yap signaling does not directly regulate these genes (Fig. 4G). In contrast, when we injected pregnant dams with tamoxifen at E17.5 to activate *Sox10-iCre^ERT2^*before birth, by P4, these glands exhibited dilated lumens marked by X-gal staining, a result of a cistronic *LacZ* reporter from the *R26^Yap5SA^*allele (Fig. 4H, black dotted line). The epithelial linings of these enlarged lumens presented nuclear Yap staining as expected (Fig. 4H, inserts), but they have lost Mist1 expression. These results support our model that heightened Yap signaling specifically disrupts acinar progenitor development during the critical prenatal period when they are separated from the basement membrane, leading to apical extrusion and apoptosis of these vulnerable cells.

### Csk-regulated tyrosine phosphorylation of Yap is dispensable for Yap function

We next asked how the loss of Csk resulted in the activation of Yap/Taz signaling. Extensive literature has documented the close relationship between Yap and SFKs ^21–24,41–45^.

Mechanistically, SFKs have been shown to directly phosphorylate Yap at multiple tyrosine residues, promoting Yap cytoplasmic-nuclear shuttling and transcriptional activity ^21–24^. Given the essential role of Csk in regulating SFKs, we examined whether Yap tyrosine phosphorylation was upregulated by *Csk* deletion. To this end, we isolated mouse embryonic fibroblast (MEF) cells from *Csk^flox/flox^*embryos and infected them with Cre-expressing adenovirus (Ad-Cre). As expected, this led to the depletion of Csk, loss of inhibitory Src^Y527^ phosphorylation and a concomitant increase in activating Src^Y416^ phosphorylation (Fig. 5A). We next performed immunoprecipitation with a pan-phosphotyrosine antibody and probed for Yap protein, revealing a strong increase in Yap tyrosine phosphorylation in the absence of Csk.

**Figure 5.**
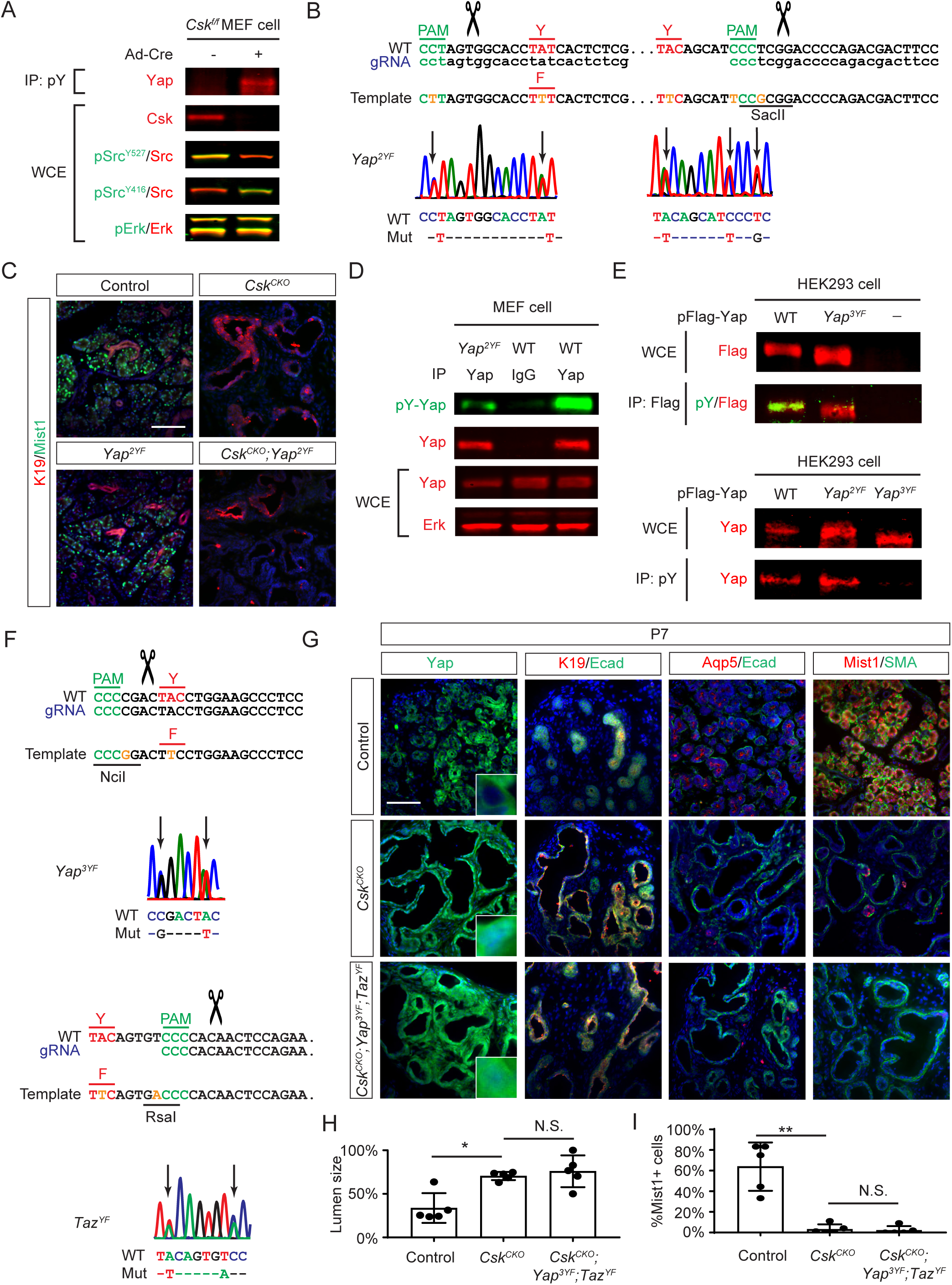
Tyrosine phosphorylation of Yap induced by *Csk* deletion does not affect Yap function. **(A)** Cre-mediated depletion of Csk in MEF cells led to the loss of pSrc^Y527^ inhibitory phosphorylation and a concomitant increase in pSrc^Y416^ activating phosphorylation without perturbing ERK phosphorylation. **(B)** CRISPR-Cas9 genome editing was used to mutate two tyrosine phosphorylation sites (*Yap^Y341F/Y357F^*or *Yap^Y2F^*). **(C)** Luminal dilation and Mist1 expression defect were not rescued in *Csk^CKO^* ;*Yap^Y2F^* mutants. **(D)** Yap tyrosine phosphorylation was reduced but not lost in *Yap^Y2F^* homozygous MEF cells. **(E)** Mutating all three tyrosine residues in a Flag-tagged Yap (Yap^Y3F^) led to a complete loss of tyrosine phosphorylation as revealed by Flag or phospho-tyrosine (pY) immunoprecipitation. **(F)** Schematic diagram of CRISPR-Cas9 genome editing to generate *Yap^Y341F/Y357F/Y394F^*(*Yap^Y3F^*) and *Taz^Y321F^* (*Taz^YF^*) mutant mice. **(G)** Yap protein remained nuclear in *Csk^CKO^* ;*Yap^Y3F^*;*Taz^YF^*mutants, resulting in luminal dilation and loss of Mist1 expression. **(H)** The lumen size was quantified as the ratio of the lumen area versus the overall epithelial area. One-way ANOVA, n=5, *P<0.01. **(I)** Quantification of Mist1+ cells. One-way ANOVA, n=5, *P<0.0001. Scale bars: 100 µm.

According to the PhosphositePlus database, the most frequently phosphorylated tyrosine residues of Yap are Y341 and Y357 (Y391 and Y407 in the Phosphosite database). We reasoned that if Csk controls Yap via Src-mediated phosphorylation, mutating these amino acids should dampen Yap activity and rescue *Csk* mutant lacrimal glands. We thus used CRISPR-Cas9 technology to generate point mutations in the *Yap* locus to replace these two tyrosine residues with phenylalanine (Fig. 5B). Surprisingly, *Yap^Y341F/Y357F^*(*Yap^2YF^*) mice were viable and fertile without any obvious phenotype. Examination of lacrimal glands showed that luminal dilation and acinar loss were also not rescued in *Csk^CKO^*;*Yap^2YF^*mutants (Fig. 5C). This prompted us to investigate whether there was significant tyrosine phosphorylation on residues other than Y341 and Y357. Indeed, western blot analysis of immunoprecipitated Yap showed that the phospho-tyrosine signal was reduced but not eliminated in *Yap^2YF^* protein (Fig. 5D). To identify the amino acid responsible for the residual tyrosine phosphorylation, we further mutated Y394 residue (Y444 in the Phosphosite database) to generate a flag-tagged *Yap^Y341F/Y357F/Y394F)^* (*Yap^3YF^*) protein. In line with a previous report, Yap^3YF^ immunoprecipitated using the anti-Flag antibody lost the phosphotyrosine signal, while the anti-phosphotyrosine antibody failed to pull down Yap^3YF^ protein (Fig. 5E) ^21^. Based on this result and the sequence homology between Yap and Taz, we used CRISPR-Cas9 technology to further generate *Yap^3YF^* mice and knock in the Y357F equivalent mutation in the *Taz* locus (*Taz^YF^*) (Fig. 5F). However, neither *Yap^3YF^* nor *Taz^YF^* mutant mice displayed any obvious abnormalities and even the introduction of both *Yap^3YF^*and *Taz^YF^* alleles into the *Csk^CKO^* background failed to prevent nuclear localization of Yap (Fig. 5G).

Accordingly, *Csk^CKO^*;*Yap^3YF^*;*Taz^YF^*mutant glands were still composed of enlarged lumens without acinar cells (Fig. 5H and I). Through exhaustive in vivo mutagenesis, we conclusively demonstrated that tyrosine phosphorylation of Yap and Taz is a fortuitous act of Src, not responsible for the regulation of Yap/Taz signaling.

### Crk and CrkL are the downstream mediators of Csk-Src signaling

If SFKs do not regulate Yap/Taz by direct tyrosine phosphorylation, there must exist intermediate factor(s) that connect the Src and Yap signaling. The DepMap consortium (depmap.org) has conducted thousands of genome-wide screens to identify the codependency of genes for cancer cell growth, providing an unbiased view of genetic relationships. We reasoned that, if we plot the codependency scores of genes shared by Src and Yap along the x and y axes, the likely intermediaries between Src and Yap would fall along a diagonal line (Fig. 6A).

**Figure 6.**
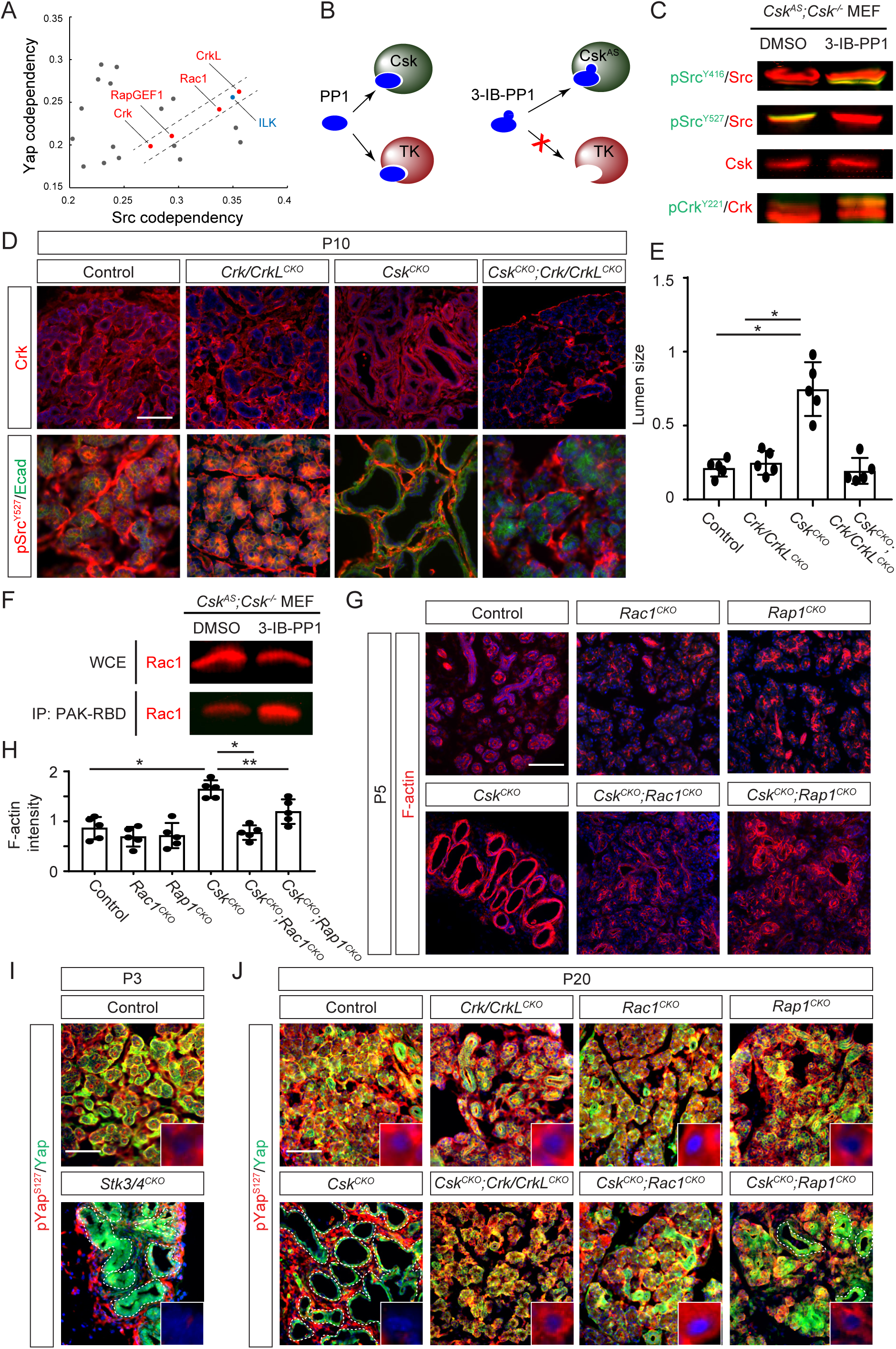
Csk-Yap signaling was mediated by the Crk/CrkL-Rac/Rap pathway. **(A)** Among the top codependency genes for both *Yap* and *Src* in Depmap data, *Crk*, *CrkL*, *Rac1* and *RapGEF1* fall along a diagonal axis. **(B)** Although PP1 is a nonselective kinase inhibitor, its bulky analog (3-IB-PP1) can only target an analog-sensitive form of Csk (Csk^AS^) that contains an enlarged ATP-binding pocket. **(C)** In MEF cells that solely express *Csk^AS^* mutant (*Csk^AS^; Csk^-/-^*), the addition of 3-IB-PP1 not only elevated pSrc^Y416^ at the expense of pSrc^Y527^ phosphorylation, but also induced Crk phosphorylation. **(D)** Deletion of *Crk*/*CrkL* did not affect pSrc^Y527^ phosphorylation, but it rescued the *Csk^CKO^* luminal dilation phenotype. **(E)** The lumen size was quantified as the ratio of the lumen area versus the overall epithelial area. One-way ANOVA, n=5, *P<0.0001. **(F)** PAK-RBD pull-down assay showed that 3-IB-PP1 enhanced the GTP-bound form of Rac1. **(G)** *Csk^CKO^* mutants exhibited a sharp increase in F-actin staining, which was diminished by the deletion of *Rac1* or *Rap1*. **(E)** Quantification of F-actin staining (arbitrary unit). One-way ANOVA, n=5, *P<0.0001. **P<0.05. **(I)** Genetic ablation of Hippo kinases encoded by *Stk3* and *4* resulted in nuclear localization of Yap, loss of pYap^S127^ phosphorylation and dilation of the lacrimal gland epithelium. **(J)** Ablation of *Crk*/*CrkL*, *Rac1* and *Rap1* in *Csk^CKO^*mutants restored pYap^S127^ phosphorylation and prevented Yap nuclear localization in acinar cells. Scale bars: 100 µm.

Remarkably, the five genes revealed by this analysis - *CrkL*, *ILK*, *Rac1*, *RapGEF1* and *Crk* - are all closely linked in integrin signaling ^46^. Specifically, it has been shown that Src phosphorylation of the scaffold protein p130Cas leads to the recruitment of Crk and its homolog CrkL, which can bind guanine exchange factors (GEFs) Dock1 and RapGEF1 to activate small GTPases Rac1 and Rap1, respectively ^47–51^. This observation places the Crk/CrkL-Rac/Rap axis as the likely candidate mediating Src-Yap signaling.

Our hypothesis predicts that the activation of Src in the absence of Csk should lead to the recruitment of Crk/CrkL and activation of Rac1 and Rap1, but this is contradicted by previous studies that failed to detect any increase in Rac1 activity in *Csk* null cells ^52,53^. We wondered whether these conflicting results may be due to the adaptation of mutant cells to the prolonged loss of *Csk*. To test this idea, we deployed a chemical genetic strategy to generate acute and specific inhibition of Csk. It exploits engineered Csk protein (Csk^AS^), which contains a gatekeeper mutation to enlarge its ATP-binding pocket, allowing it to accommodate a bulky 3-iodo-benzyl group (3-IB) attached to a highly potent but nonselective kinase inhibitor PP1 (Fig. 6B) ^54^. On the other hand, the 3-IB modification prevents this PP1 derivative from locking into the conserved catalytic domains, rendering it ineffective against other endogenous protein kinases. When expressed from a bacterial artificial chromosome (BAC) transgene to ensure the endogenous expression pattern, *Csk^AS^* is able to support normal mouse development in *Csk^AS^;Csk^-/-^* animals ^54^, demonstrating that Csk^AS^ can functionally substitute for the wild type Csk. We confirmed that MEF cells isolated from *Csk^AS^;Csk^-/-^*animals were susceptible to acute inhibition of Csk signaling by 3-IB-PP1, which caused a rapid reduction in pSrc^Y527^ and an increase in pSrc^Y416^ phosphorylation without altering the level of Csk protein (Fig. 6C).

Importantly, treatment with 3-IB-PP1 also led to a significant increase in pCrk^Y221^, a known marker of Crk activity. These results showed that Csk is indeed an upstream regulator of Crk function.

To investigate whether the Crk family protein participates in Csk signaling in lacrimal gland development, we next performed a genetic rescue experiment by deleting both *Crk* and *CrkL*. Although immunostaining showed that Crk was efficiently ablated in the epithelium in *Crk/CrkL^CKO^* mutants, the lacrimal gland morphology was unaffected. However, in stark contrast to *Csk^CKO^* mutants, the size of the lumens in P10 *Csk^CKO^;Crk/CrkL^CKO^* lacrimal glands was reverted to normal despite the persistent loss of pSrc^Y527^ phosphorylation (Fig. 6C and D). This demonstrated that *Crk* and *CrkL* are essential for *Csk* mutant phenotypes.

### Rac1 and Rap1 suppress the Hippo-Yap pathway in response to Csk-Src signaling

Encouraged by the genetic rescue of the *Csk* mutant by the deletion of *Crk*/*CrkL*, we next pursued their downstream small GTPases. Notably different from previous studies in *Csk^-/-^* cells ^52,53^, 3-IB-PP1 treatment of *Csk^AS^;Csk^-/-^*MEF cells led to a significant increase in the level of the GTP-bound Rac1 detected by pull-down with the RBD domain of PAK (PAK-RBD) (Fig. 6F). This showed that acute inhibition of Csk indeed led to the activation of Rac1. Consistent with the role of Rac1 in promoting actin polymerization, the *Csk^CKO^* mutant lacrimal gland epithelium also exhibited a significant increase in F-actin staining, which was reversed by the deletion of *Rac1* (Fig. 6G and H). Importantly, the lumen dilation phenotype was also ameliorated in *Csk^CKO^;Rac1^CKO^*glands. On the other hand, deletion of the two *Rap1* genes, *Rap1a* and *Rap1b*, in *Csk^CKO^;Rap1^CKO^*animals similarly reduced F-actin staining and partially normalized lacrimal gland architecture. These results suggested that Rac1 and Rap1 are important mediators of Csk signaling.

The actin cytoskeleton network regulated by small GTPases plays an important role in Hippo signaling, whose mammalian orthologs Mst1/2 (encoded by *Stk3/4*) activate Lats1/2 kinases to prevent Yap nuclear translocation by phosphorylating multiple serine residues of Yap, especially Yap^S127^ ^30,55^. We thus explored the consequence of impaired Hippo signaling by ablating *Stk3/4* in the lacrimal gland epithelium, which showed greatly reduced pYap^S127^ staining and nuclear localization of Yap as expected. Importantly, *Stk3/4^CKO^* displayed enlarged lumens like *Csk^CKO^* mutants, demonstrating that the loss of Hippo signaling was sufficient to reproduce the *Csk* phenotype (Fig. 6I, dotted lines and inserts). Further supporting the view that the deletion of *Csk* may disrupt Hippo signaling, pYap^S127^ staining was abundant in the cytoplasm of control lacrimal gland cells, but it was completely lost in the *Csk^CKO^* mutant epithelium, which is the exact opposite pattern compared to Yap nuclear staining (Fig. 6J, dotted lines and inserts).

Although the normal pattern of pYap^S127^ was not affected by the deletion of *Crk*/*CrkL*, *Rac1* and *Rap1* alone, it was restored in *Csk^CKO^;Crk/CrkL^CKO^* and *Csk^CKO^;Rac1^CKO^*acinar cells. In the partially rescued *Csk^CKO^;Rap1^CKO^* mutants, the remaining dilated lumens inevitably lost pYap^S127^ staining while retaining nuclear Yap expression. These results indicate that deletion of *Crk/CrkL*, *Rac1* and *Rap1* suppressed Yap activity in the lacrimal gland by maintaining Hippo signaling.

Lastly, we showed that the lumen size was normalized in *Csk^CKO^;Crk/CrkL^CKO^* , *Csk^CKO^;Rac1^CKO^* and, to a lesser extent, *Csk^CKO^;Rap1^CKO^*mutants. In addition, the pattern of SMA+ myoepithelial cells and Krt19+ ducts and the quantity of Mist1/Aqp5-expressing acinar cells were similarly recovered (Fig. 7A and B). Taken together, our study demonstrated that Csk-Src signaling is transmitted via the Crk/CrkL-Rac/Rap cascade to regulate Hippo-Yap signaling in lacrimal gland development.

**Figure 7.**
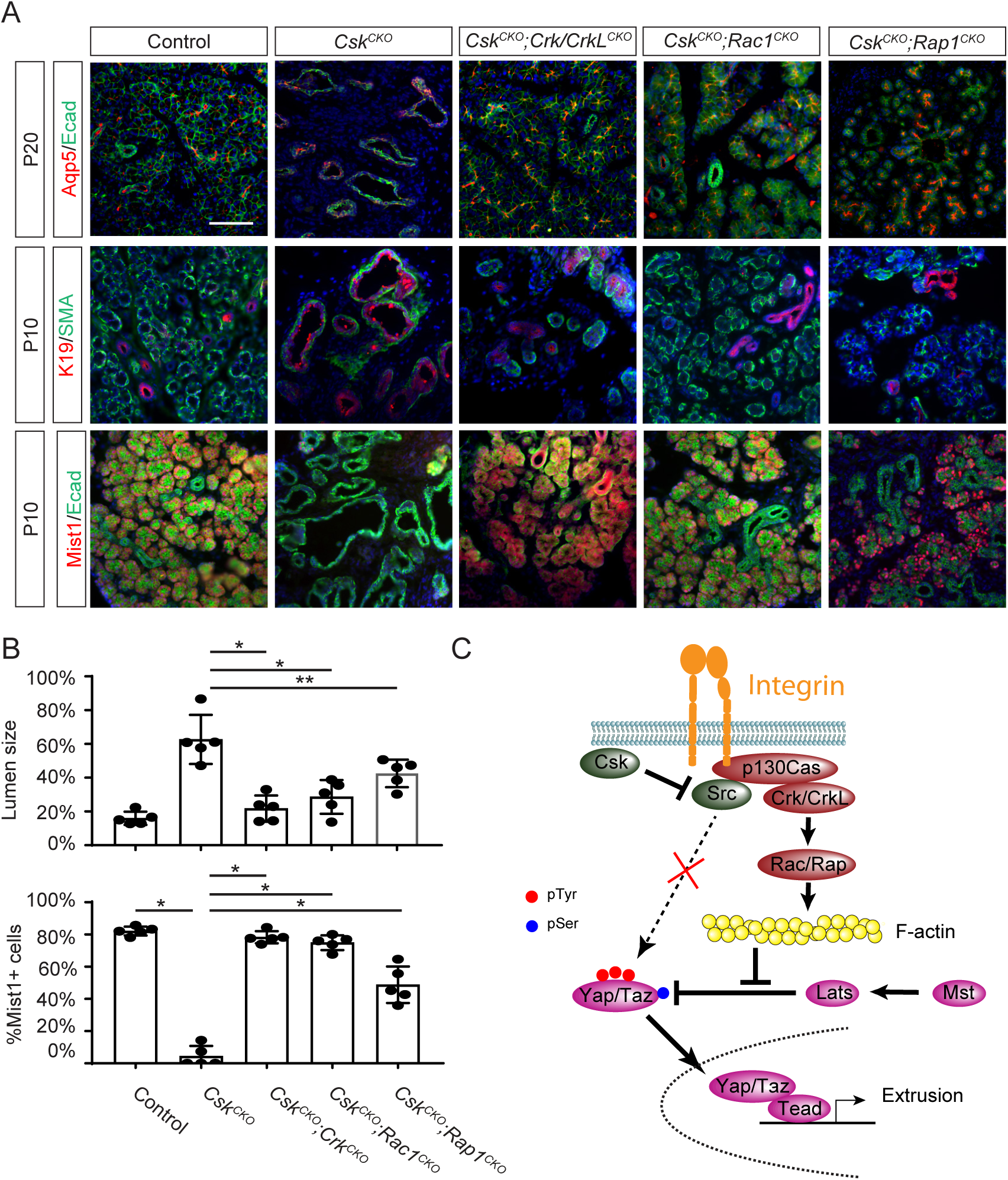
Deletion of *Crk*/*CrkL*, *Rac1* or *Rap1* rescued *Csk^CKO^* mutant lacrimal gland phenotype. **(A)** The normal glandular morphology and Mist1-expressing acini were restored after the genetic deletion of *Crk*/*CrkL*, *Rac1* or *Rap1* in *Csk^CKO^* mutants. **(B)** Quantification of the lumen size and the number of Mist1+ cells. One-way ANOVA, n=5, *P<0.0001, **P<0.05. **(C)** The model of Csk control of Yap signaling. Csk-regulated Src kinase phosphorylates p130Cas in the Integrin-anchored focal adhesion complex to create docking sites for Crk and CrkL, which recruit GEFs for small GTPases Rac and Rap to promote F-actin formation. This in turn suppresses serine phosphorylation (pSer) of Yap by Mst-Lats signaling, leading to nuclear translocation of Yap which promotes cell extrusion and abolishes acinar differentiation. In contrast, fortuitous tyrosine phosphorylation of Yap by Src is dispensable for Yap activity. Scale bar: 100 µm.

## Discussion

In the current study, we have uncovered a previously unappreciated requirement for regulating cell extrusion in lacrimal gland development. We showed that the deletion of *Csk* in the lacrimal gland epithelium resulted in luminal dilation and the loss of acinar cells, which originated from apoptosis of lacrimal gland progenitors caused by Src-induced cell extrusion. Through genetic analysis, we challenged the widely held belief that Src directly regulates Yap via tyrosine phosphorylation. Instead, considering the known role of Src in regulating integrin signaling by phosphorylating the p130Cas protein ^1^, we propose that the main function of Src is to create docking sites on p130Cas for Crk and CrkL that subsequently activate Rac and Rap to promote F-actin polymerization (Fig. 7C). This prevents Hippo signaling from suppressing the nuclear translocation of Yap and Taz to stimulate Tead transcription factors, causing expulsion of acinar progenitor cells into the lacrimal gland lumen. Our study illuminated the molecular mechanism by which Src-regulated mechanosignaling induces cell extrusion by Yap/Taz-mediated transcriptomic reprogramming.

Cell extrusion is a crucial process in development, tissue homeostasis and tumor suppression ^56,57^. It is a key element of the cell competition program that removes unfit cells, where mutant clones are actively eliminated by surrounding wild type, but not by other mutant cells ^58,59^. This is exemplified in *Drosophila*, where *Csk* knockdown in the entire eye imaginal discs resulted in organ overgrowth, but discrete inactivation of *Csk* in clonal patches led to epithelial extrusion and apoptotic death ^5^. In contrast, we found that deleting Csk throughout the lacrimal gland epithelium specifically triggers the loss of acinar cells, which extruded apically into the lacrimal gland lumen – a stark difference from the basal extrusion observed in Drosophila. To explain these discrepancies, we note that the acinar progenitors are unique among the lacrimal gland cells in their separation from the basement membrane by myoepithelial progenitors during embryonic development. Unencumbered by the integrin-mediated adhesion to the basement membrane, these acinar progenitors are receptive to mobilization by Src-activated Yap signaling to delaminate from the lacrimal gland epithelium. Supporting this model, we observed loss of acinar cells only when Yap was activated in prenatal progenitors, but not in mature lacrimal glands, coinciding with the critical period when acinar progenitors lack basement membrane anchorage. It is intriguing to note that Src is known to act in the integrin pathway to promote cell invasion and mobility by stimulating the disassembly and turnover of the focal adhesion complex. In the absence of active integrin engagement with the basement membrane, the deletion of Csk may have unleashed Src to hijack the cell motility component of the integrin pathway to promote cell extrusion. This study highlights that cell extrusion can arise not just from competition between different genotypes, but also from differences in adhesive properties and motility potential among cells harboring the same mutations.

Comprising nine members and present in all cell types, SFKs exert a profound influence across the proteome, as evidenced by the lethal outcomes of their genetic knockouts and their potency for driving oncogenic transformation ^1^. Although mass spectrometry analyses have shown that SFKs may phosphorylate hundreds of proteins, identifying their physiologically relevant targets remains an ongoing challenge ^60,61^. In the absence of Csk, unfettered SFK signaling could have pleiotropic effects through numerous downstream effectors. For example, in *Drosophila*, the extrusion of *Csk*-deficient cells in the eye is thought to be mediated by the Cadherin-JNK-MMP axis ^5^. In contrast, in MDCK cells, a standard mammalian model for studying cell extrusion, the process in Src- and Yap-activated cells was reported to rely on MAPK and PI3K-mTOR signaling pathways, respectively ^6,62^. It is thus remarkable that the striking lacrimal gland phenotypes in *Csk* mutants can be completely replicated by activating Yap and reversed by its deletion. This finding suggests that at least in the context of lacrimal gland development, the transcriptomic changes driven by Yap are both necessary and sufficient to mimic the broad phosphoproteomic changes caused by SFKs. Our study highlights a critical connection between these two proto-oncoproteins, potentially offering insights applicable to both developmental and cancer biology.

Yap was originally named as “<u>Y</u>es-<u>a</u>ssociated <u>p</u>rotein” because of its binding affinity to the SFK family member Yes ^63^, but the biological significance of this association remains contentious in the field ^64^. Previous studies have posited that Src phosphorylation of Yap may control its nuclear localization ^21^ or transcriptional activity ^22–24^, but these assertions have been disputed by other research suggesting that Src regulates Yap indirectly via PI3K, JNK or tyrosine phosphorylation of Lats ^41–44^. Similar controversy also exists concerning Src’s regulation of Taz ^65,66^. These discrepancies likely arise because these studies have relied on gene overexpression, RNAi knockdown or small molecule inhibitors, which are prone to off-target artifacts and may not accurately reflect physiological conditions. Most importantly, these models are all based on in vitro cell culture systems, but were not tested rigorously in vivo. Our work helps to fill this gap by generating knock-in mice to systematically mutate tyrosine residues phosphorylated by Src in Yap and Taz. Contradicting the prevailing belief that tyrosine phosphorylation is crucial for Yap/Taz regulation, these phosphorylation-deficient animals didn’t exhibit any obvious defects in growth and fertility, nor did they rescue lacrimal gland development in the *Csk* mutant background. These findings lead us to conclude that, despite being observed in numerous experimental settings, tyrosine phosphorylation of Yap/Taz is not physiologically relevant to Yap signaling. Instead, we showed that the *Csk* lacrimal gland phenotype can be mimicked by deactivating Hippo kinases Mst1/2 and rescued by genetically ablating *Crk*/*CrkL*, *Rac* and *Rap*, which reinstates Hippo-dependent phosphorylation and cytoplasmic localization of Yap. Our mouse genetic study breaks new ground by revealing a previously unrecognized role of Crk proteins in Yap regulation, establishing a Src-Crk/CrkL-Rac/Rap molecular pathway that links Csk to Hippo signaling.

## Methods and Materials

### Mice

All procedures related to animal care and experimentation were conducted in adherence to the protocols and guidelines approved by the Institutional Animal Care and Use Committee at Columbia University. Mice carrying *Crk^flox^, CrkL^flox^*, *Krt7^Cre-ERT2^*, *R26^Yap5SA^, Rap1a^flox^*and *Rap1b^flox^* alleles were bred and genotyped as outlined ^36,38,51,67,68^. *Csk^flox^* is from Dr. Alexander Tarakhovsky (The Rockefeller University, New York, NY) ^69^, *Le-Cre* from Richard Lang (Children’s Hospital Research Foundation, Cincinnati, OH) ^70^, *Mist1^Cre-ERT2^* from Timothy Wang (Columbia University, New York, NY) ^34^, *SMA-Cre^ERT2^* from Ivo Kalajzic (University of Connecticut, Farmington, CT) ^35^, *Rac1^flox^* from Dr. Feng-Chun Yang (Indiana University School of Medicine, Indianapolis, IN) ^71^, and *Ai9* (Stock #: 007908), *Csk^AS^;Csk^-/-^* (Stock #: 025250), *R26R* (Stock #: 003474), *Sox10-iCre^ERT2^* (Stock #: 027651), *Stk3 ^flox^;Stk4^flox^* (Stock #: 017635) and *Yap^flox^;Taz^flox^* (Stock#: 030532) from Jackson Lab. Mice were maintained on a mixed genetic background, and at least three animals were analyzed for each described cross, using Cre-heterozygous mice as controls. Tamoxifen, prepared in corn oil at 10 mg/mL and stored at 4°C, was administered via intraperitoneal injections at 100 mg/g of mouse body weight, with three consecutive injections performed to achieve maximum Cre induction.

*Yap^2YF^*, *Yap^3YF^* and *Taz^YF^*mice were generated using the modified *Easi*-CRISPR method ^72^. Briefly, highly stringent gRNAs near the mutation sites were identified using online design tools (http://crispor.tefor.net) and chemically synthesized by IDT with 2’-O-methyl 3’phosphorothioate modifications and end-blocking Alt-R modifications. Single-stranded donor templates, with the desired amino acid alterations and disrupted PAM sites, along with silent mutations creating restriction enzyme sites, were injected into mouse zygotes with a pre-assembled gRNA-Cas9 enzyme complex by the transgenic facility at Columbia University Medical Center. Correctly targeted founder animals were identified through direct sequencing. The following gRNAs were used: GGAAGTCGTCTGGGGTCCGA GGG and CGAGAGTGATAGGTGCCACT AGG for *Yap^2YF^*, GGAGGGCTTCCAGGTAGTCG GGG for *Yap^3YF^* and GGAAGTCTTCTGGAGTTGTG GGG for *Taz^YF^*.

### Histology and immunohistochemistry

Histology and immunohistochemistry were performed on the paraffin and cryosections as previously described ^73,74^. For hematoxylin and eosin (H&E) staining, 10 µM paraffin sections underwent deparaffinization with histosol wash, rehydration through decreasing concentrations of ethanol solutions, and final washing in water. The slides were immersed in hematoxylin for 3 minutes, followed by a 10-15 minute wash with tap water. Subsequently, they were decolorized with 1% acid alcohol for 30 seconds before treatment with eosin for 1 minute. Samples were dehydrated through increasing ethanol concentrations, transferred to histosol, and mounted using Permount mounting medium.

For immunofluorescence staining, cryosection slides were briefly washed with PBS to remove OCT. Antigen retrieval was performed in a microwave by boiling for 1-2 minutes, followed by heating for 10 minutes at low-power settings in sodium citrate buffer (10 mM sodium citrate, pH 6.0). Sections were then washed with PBS and blocked for 30 minutes with 5% normal goat serum (NGS)/0.1% Triton in PBS. Primary antibody incubation occurred overnight at 4°C in a humid chamber, followed by incubation with fluorophore-conjugated secondary antibodies for 2 hours at room temperature in the dark.

TUNEL assays were carried out on 10 µm cryosections following the manufacturer’s protocol in the Fluorescein in Situ Cell Death Detection kit (Roche Applied Science, Indianapolis, IN).

The following primary antibodies were utilized: Aquaporin5 (abcam #ab78486), Cleaved caspase3 (CST#9662), Col2A1 (Abcam #ab34712), Crk (BD #610036), Csk (Proteintech #17720-AP), c-Src (Santa Cruz #sc-8056), E-cadherin (BD Bio #610181 and CST #3195), Keratin14 (Covance #PRB-155P), Keratin19 (DSHB #TROMA-III), Ki67 (BD #550609), Laminin (BD #354232), Mist1 (abcam #ab187978), p63 (Biolegend #619001), Pax6 (Covance #PRB-278P), Phalloidin (Life Technology #A12380), pHH3 (Millipore #06-570), pSRC527 (CST #2105), pYAP Ser127 (CST #4911), SMA (Agilent #M085129-2), Yap (CST #14074 and Santa Cruz #sc-101199), Yap-taz (CST #8418), ZO1 (DSHB #R26.4C). All commercial antibodies were validated by vendors. At least three embryos of each genotype were stained for each marker.

### Single-cell RNA-seq and bioinformatic analysis

Lacrimal glands obtained from P1 pups (three for each genotype) underwent a PBS wash for 2-5 minutes and were digested with dispase (2000 units/ml) and papain (20 units/ml) for 30 minutes at room temperature. After gently removing the digestion enzyme mixture, the glands were washed with PBS and mechanically disrupted through vigorous pipetting until single-cell dispersion was observed under the microscope. Following three washes in PBS and centrifugation at 3000 rpm at 4°C, control and *Csk^CKO^*mutant cells were treated with cell multiplexing oligos (CMO) in accordance with the 10x Genomics 3’ cellplex kit manual. The cells were then pooled together in a single collection tube for library preparation and subsequent sequencing at Columbia University’s single-cell sequencing core. The RNAseq data were deposited at the Gene Expression Omnibus (GEO) database (GSE253784) and single-cell portal (SCP2499).

Gene-cell matrices, generated using Cell Ranger 2.1.1 from 10x Genomics, were imported into R and analyzed using Seurat. Cells with <200 or > 4000 genes, as well as those exhibiting >15% expression of mitochondrial genes, were filtered out prior to dataset normalization and scaling. Dimensionality reduction was achieved through Principal Component Analysis (PCA) and non-linearly with Uniform Manifold Approximation and Projection (UMAP) with dimensions = 1:20. Following the integration of control and mutant datasets through canonical correlation analysis (CCA), clustering was performed using “RunUMAP” and annotated based on known marker genes. Differential gene expression testing was conducted using the “FindMarkers” and “FindAllMarkers” functions in Seurat.

A subset comprising epithelial clusters and the Mes6 cluster was extracted and re-clustered. A diffusion map of this subset was generated using the R package “destiny” ^28^. Trajectory analysis and pseudotime analysis for this subset were performed using the Monocle package in R ^37^. Gene Set Enrichment Analysis (GSEA) of hallmark pathways between control and mutant samples was calculated using the R package “fgsea.” Regulon analysis was executed using the pySCENIC packages in Python ^37^.

### Cell culture, immunoprecipitation and western blot

Mouse embryonic fibroblast (MEF) cells were isolated from E13.5 mouse embryos. Following the removal of heads and guts with autoclaved scissors, cleaned embryos were finely chopped and incubated with trypsin for 30 minutes at 37°C. Further dissociation was achieved through vigorous pipetting, and single cells were cultured in a 10 cm cell culture dish with DMEM media containing 10% FBS. *Csk^AS^;Csk^-/-^*MEF cells were treated with 25 µM 3IB-PP1 for 5 minutes, immediately washed with ice-cold PBS, and resuspended in lysis buffer for subsequent analysis. The Rac1-GTPase assay was performed using the Rac1 Pull-Down Activation Assay Biochem Kit (Cytoskeleton #BK-035).

*Csk^flox/flox^* MEF cells, at 60% confluency in a 15 mm dish, were incubated with Ad5-CMVCre virus (1x10^10^ pfu/ml, Gene Transfer Vector Core, University of Iowa, IA) at a 1:500 dilution for 5 days before being treated with 1 mM sodium orthovanadate (S6508, Sigma Aldrich) for 3 minutes at 37°C. After adding ice-cold lysis buffer (C2978, Sigma Aldrich) containing 1X protease inhibitor and 1 µM sodium orthovanadate, cells were scraped from the culture plate, collected in a 1.5 ml tube, and kept on ice for 10 minutes. Following dissociation by passing through syringe needles ten times and sonication three times with a 5-second pulse each, the cell lysate was centrifuged at 15,000 rpm for 10 minutes. The supernatant was incubated with immunoprecipitation antibodies at the suggested concentration overnight at 4°C, followed by protein A/G agarose beads (Santa Cruz Biotechnology, SC2003) for 2 hours at 4°C. The beads were collected by centrifugation at 4°C at 5000 rpm for 2 minutes and washed with ice-cold PBS in the presence of 1X protease inhibitor and 1 mM sodium orthovanadate three times for 5 minutes each. After the beads were resuspended in 40 µl of 1X Laemmli buffer and boiled at 100°C for 5 minutes, 20 µl of the supernatant was loaded onto a 12% SDS PAGE agarose gel for standard western blot analysis. Antibodies against phospho-Tyrosine (P-Tyr-100) (CST #9411), YAP (CST #14074 and Santa Cruz #sc-271134), and Flag (CST # 8146) were used for immunoprecipitation (5 µg antibody for 500 µg protein). Antibodies against Crk (BD #610036), Csk (Proteintech #17720-AP), ERK1/2 (CST #4695), pCRK (CST #3491), pERK1/2 (Santa Cruz Biotechnology, sc-7383), pSRC416 (CST #6943), pSRC527 (CST #2105), YAP (CST #14074), and phospho-tyrosine (P-Tyr-100) #9411 were used in western blot (1:1000 dilution).

### Cell transfection

The plasmid encoding wild-type Yap was obtained from Addgene (pcDNA Flag Yap1, Addgene #18881). Plasmids encoding *Yap^2YF^* and *Yap^3YF^* were constructed from pcDNA Flag-Yap1 Y357F (Addgene #18882) through site-directed mutagenesis using Neb Q5 high-fidelity polymerase (New England Biolab #M0491). The primers used for Y349F were Forward: CAGTGGCACCTATCACTCTCGAGATGAG and Reverse: TTAAGGAAAGGATCTGAGCTATTG. The primers used for Y402F were Forward: TTTCCCAGACTTCCTTGAAGCCATTC and Reverse: CGGTTCTGCTGTGAGGGC.

For cell transfection, 10 µg of plasmid and 30 µg of polyethylenimine (PEI, Serochem #Prime p100) were separately incubated with 300 µl of DMEM media each for 10 minutes at room temperature. They were then mixed and gently added to 70% confluent HEK293 cells in a 10 cm culture dish with 10 ml DMEM media. After overnight incubation, the culture media was replaced with DMEM containing 10% FBS, and the transfected cells were harvested the following day.

### Quantification and Statistical Analysis

Lumen sizes were quantified as the area ratios of the inner lumens versus the outer epithelium linings using Image J. For the quantification of F-actin staining, fluorescent images were converted to grayscale, and mean intensities were measured using ImageJ. To quantify Mist1 expression, cell proliferation, and apoptosis, the percentage of Mist1, Ki67, and TUNEL-positive cells were normalized against the total number of DAPI-positive cells within each epithelial cluster. At least three animals for each genotype were analyzed. Statistical analysis was performed using GraphPad Prism 7. Sample sizes were not predetermined. Data represent mean ± s.d. An unpaired two-tailed t-test was used for comparing two conditions and one-way ANOVA with Tukey’s multi-comparison test for three or more conditions.

### Database Analysis

The tyrosine phosphorylation sites in Yap1 were identified from the PhosphoSitePlus database (phosphosite.org) by examining the number of references that uncovered phosphorylation events in each tyrosine residue.

The top codependencies for *Yap1* and *Src* were retrieved from the Depmap database (Public 23Q4+Score, Chronos) at depmap.org, and plotted in a 2-D chart. Genes with similar codependency scores to both *Yap1* and *Src* were identified as lying within two diagonal lines bisecting the plot.

## Acknowledgements

The authors thank Drs. Ruth Ashery-Padan, Tom Curran, Ivo Kalajzic, Richard Lang, Lawrence A Quilliam, Feng-Chun Yang, Alexander Tarakhovsky and Timothy Wang for mice and reagent. The work was supported by grants from NIH (R01EY018868 and R01EY031210 to X.Z. and R01HL152293 to J.Q.). Q.W. is supported by a Pathway to Independence Award (K99EY032171) and N.M. is supported by a Knights Templar Eye Foundation Career Starter Grant Award. The Columbia Ophthalmology Core Facility is supported by NIH Core grant 5P30EY019007 and unrestricted funds from Research to Prevent Blindness (RPB).

## Author contributions

A.H., Q.W., Y.W., N.M. and X.Q. performed experiments. A.H., Q.W., Y.W and X.Z. analyzed data. A.H., J.M., JQ, WC, and X.Z. wrote and edited the manuscript.

---

**Supplementary Figure 1.**
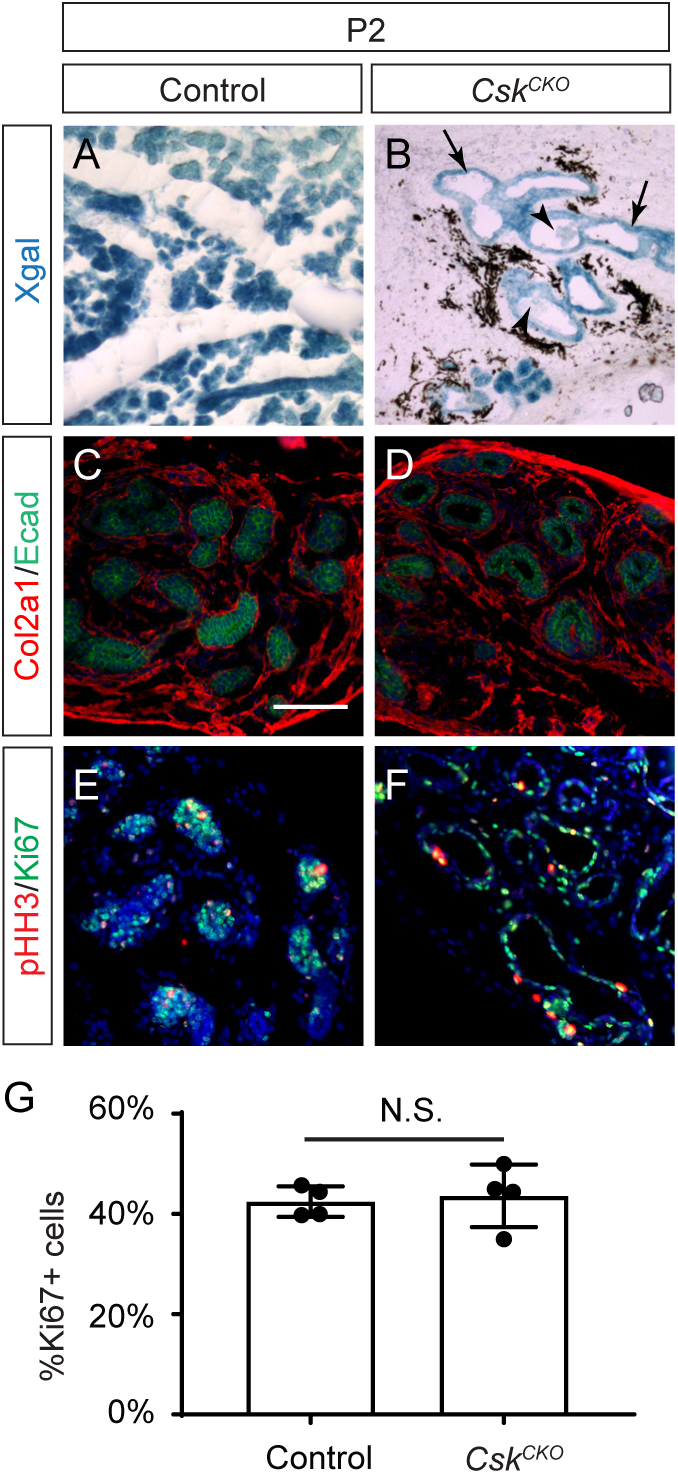
The epithelial-specific deletion of *Csk* did not affect mesenchymal differentiation or epithelial cell proliferation. **(A-B)** The *R26R* reporter activity (Xgal staining) was restricted to the *Csk^CKO^* mutant epithelium. **(C-D)** The mesenchymal expression of Col2a1 was unaffected in *Csk^CKO^* mutants. **(C-D)** The staining of Ki67 and pHH3 showed no changes in cell proliferation between control and *Csk^CKO^* mutant lacrimal glands. **(G)** Quantification of Ki67+ cells. Student’s t test, n=4, N.S. not significant. Scale bar: 100 µm.

**Supplementary Figure 2.**
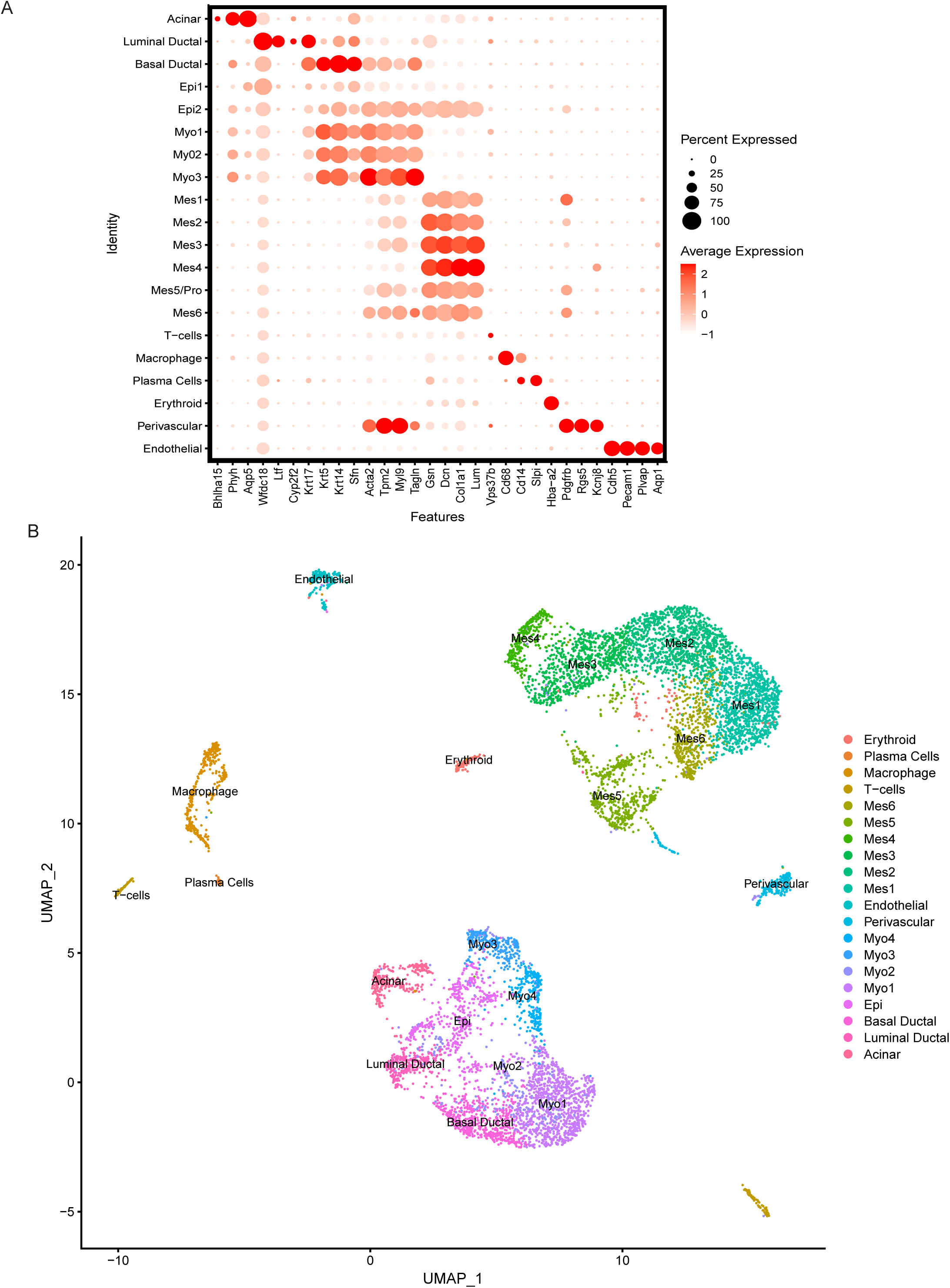
Single-cell RNAse analysis of *Csk^CKO^* mutants. **(A)** Dot plot of cell type specific markers used to annotate the integrated single-cell RNAseq data. **(B)** Clustering of the integrated single-cell RNAseq data depicted in UMAP.

**Supplementary Figure 3.**
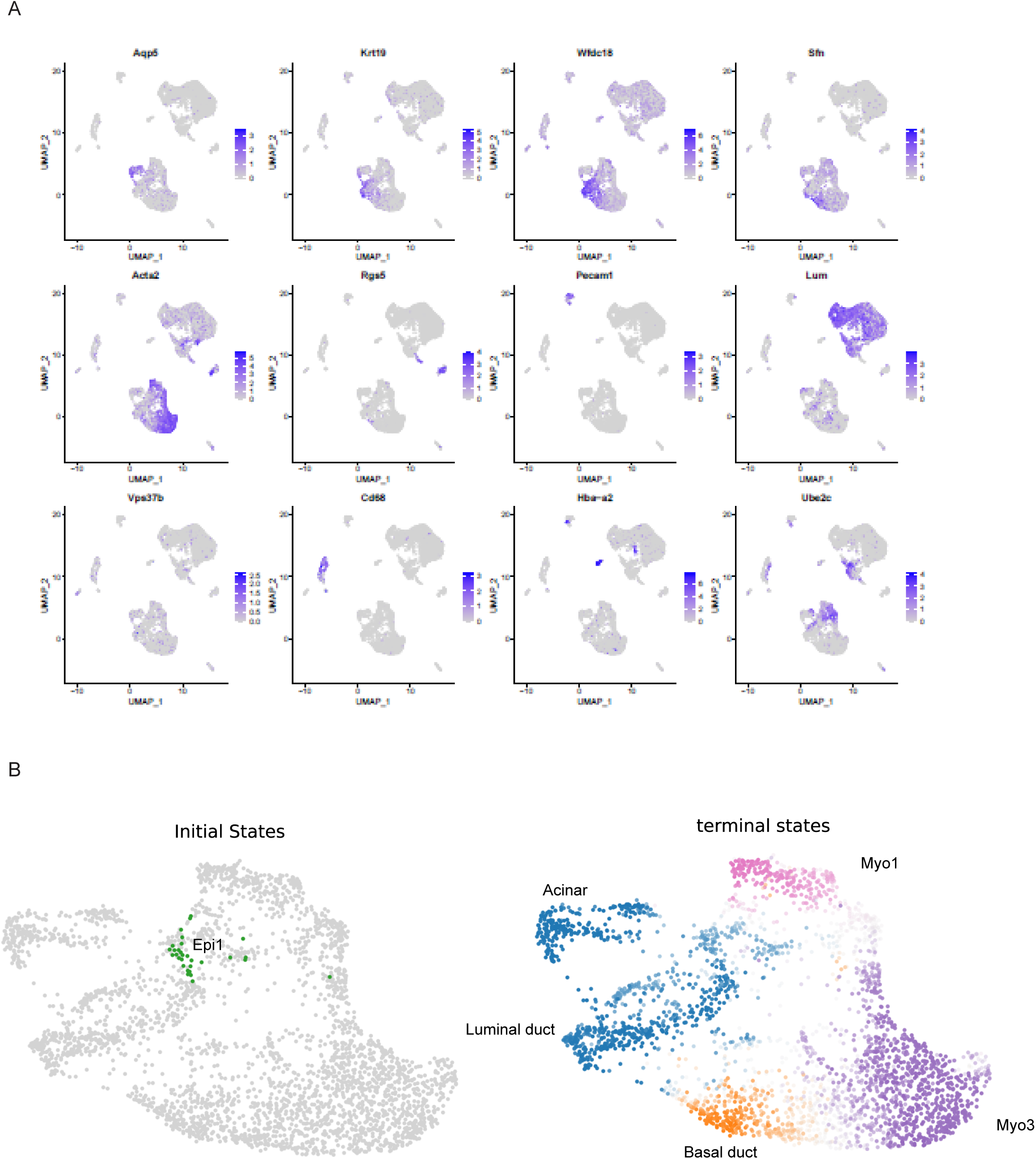
Feature plots and RNA velocity analysis of single-cell RNAse data. **(A)** Feature plots of markers used to annotate single-cell RNAseq data. **(B)** RNA velocity analysis identified the Epi1 cluster at the initial state of the epithelial lineage, whereas the myoepithelial, duct and acinar clusters are the terminal states.

